# A modeling framework for adaptive lifelong learning with transfer and savings through gating in the prefrontal cortex

**DOI:** 10.1101/2020.03.11.984757

**Authors:** Ben Tsuda, Kay M. Tye, Hava T. Siegelmann, Terrence J. Sejnowski

## Abstract

The prefrontal cortex encodes and stores numerous, often disparate, schemas and flexibly switches between them. Recent research on artificial neural networks trained by reinforcement learning has made it possible to model fundamental processes underlying schema encoding and storage. Yet how the brain is able to create new schemas while preserving and utilizing old schemas remains unclear. Here we propose a simple neural network framework based on a modification of the mixture of experts architecture to model the prefrontal cortex’s ability to flexibly encode and use multiple disparate schemas. We show how incorporation of gating naturally leads to transfer learning and robust memory savings. We then show how phenotypic impairments observed in patients with prefrontal damage are mimicked by lesions of our network. Our architecture, which we call DynaMoE, provides a fundamental framework for how the prefrontal cortex may handle the abundance of schemas necessary to navigate the real world.

## Introduction

Humans and animals have evolved the ability to flexibly and dynamically adapt their behavior to suit the relevant task at hand [1]. During a soccer match, at one end of the pitch a player attempts to stop the ball from entering the net. A few moments later at the opposite end of the pitch, the same player now tries to put the ball precisely into the net. To an uninitiated viewer, such apparently contradictory behaviors in nearly identical settings may seem puzzling, yet the ease with which the player switches between these behaviors (keep ball away from net or put ball into net) highlights the ease with which we adapt our behavior to the ever-changing contexts (near own net or opposing team’s net) we experience in the world. A bulk of evidence from observations of humans with prefrontal cortical lesions, neuroimaging studies, and animal experiments have indicated the importance of the prefrontal cortex (PFC) and connected regions in encoding, storing, and utilizing such context-dependent behavioral strategies, often referred to as mental schemas [2],[3],[4],[5]. Yet how the prefrontal and related areas are able to translate series of experiences in the world into coherent mental schemas which can then be used to navigate the world remains unknown.

Research in reinforcement learning has helped provide some insight into how the PFC may transform experiences into operational schemas [6],[7],[8]. In reinforcement learning paradigms, an agent learns through trial and error, taking actions in the world and receiving feedback [9]. Recent work has demonstrated how recurrent neural networks (RNNs) trained by trial-by-trial reinforcement learning can result in powerful function approximators that mimic the complex behavior of animals in experimental studies [8].

Although reinforcement learning has provided invaluable insight into the mechanism the PFC may use, it remains unclear how the PFC is able to encode multiple schemas, building on each other, without interference, and persisting so they may be accessed again in the future. The majority of models capable of solving multi-strategy problems require specially curated training regimens, most often by interleaving examples of different problem types [10]. Models learn successfully due to the balanced presentation of examples in training; if the training regimen is altered—for example, problem types appear in sequence rather than interleaved, as often happens in the world—the unbalanced models fail miserably [11].

Some techniques have been proposed to help models learn and remember more robustly, yet none have established how these processes may occur together in the brain. For example, continual learning techniques (e.g. [12],[13]) propose selective protection of weights. Yet such techniques heavily bias networks toward internal structures that favor earlier, older experiences over newer ones and are potentially not biologically realistic [10]. Other models either require explicit storage of past episodes for constant reference [14],[15], or an “oracle” to indicate when tasks are “new” [16],[17].

We propose a framework for how the PFC may learn through reinforcement and adapt to new environments without “oracle” supervision, while remaining robust against catastrophic forgetting. Our model adopts a hierarchical gating structure for the PFC that mirrors the mixture of experts (MoE) class of models [18] (Fig. 1a) widely found in machine learning applications [19],[20],[21],[22]. We demonstrate how such a hierarchical gated architecture naturally leads to transfer learning as new scenarios are encountered. Furthermore, we show how our network adaptively learns and, due to its architecture, demonstrates robust memory savings for past experiences. Our framework provides a basis for how the PFC and related areas may encode, store, and access multiple schemas.

**Fig.1.**
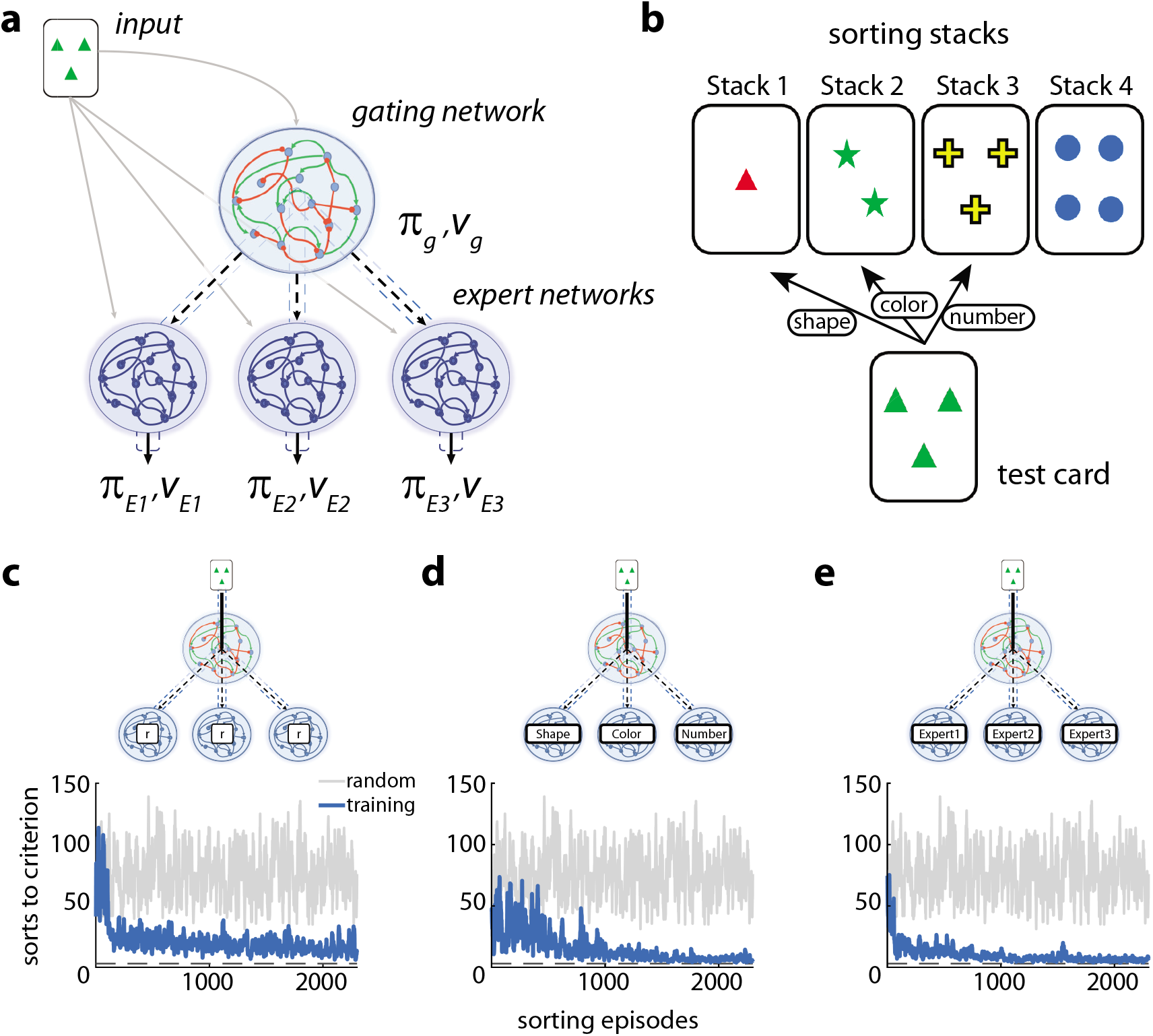
DynaMoE network structure and the WCST. **a**, The DynaMoE network is in the MoE family of networks. A gating network takes input and outputs a decision of which expert network to use (*π_g_*) and a value estimate (*v_g_*). The chosen expert network (e.g. *E*_1_) takes input and outputs an action to take (*π*_*E*1_)—for the WCST, in which stack to place the current card—and a value estimate (*v*_*E*1_). **b**, The WCST. The subject must sort the presented test card to one of four stacks by matching the relevant sort rule. The subject continues to attempt sorting test cards until achieving the termination criterion of sorting a given number of cards correctly in a row. **c–e** MoE networks on the classic WCST. **c**, MoE network with 3 experts achieves good performance quickly and slowly improves further over time. **d**, MoE network with pretrained experts on the sort rules also learns quickly reaching near perfect performance faster. **e**, DynaMoE network pretrained sequentially on the sort rules learns rapidly and reaches near perfect performance fastest. In all plots, blue traces are from networks during training and grey traces are random behavior for reference. Grey dotted line is the minimum sorts to criterion.

## Results

To demonstrate the properties of our framework we chose the Wisconsin Card Sorting Task (WCST), a commonly used clinical assessment of PFC function [2],[5],[23]. In the WCST, a subject is required to sequentially sort cards according to one of three possible sorting rules: shape, color, number (Fig. 1b; see Methods for full description). The sorting rule is not explicitly given, but rather must be discovered through trial and error. After each attempted card sort, the subject receives feedback as to whether the sort was correct or incorrect. After a set number of correct card sorts in a row, the sort rule is changed without signal, requiring the subject to adapt behavior accordingly [5],[23]. Performance can be measured by the number of attempted card sorts until the episode termination criterion is achieved (“sorts to criterion;” 3 correct sorts in a row in our simulations), with fewer attempted sorts representing superior performance.

The WCST requires the PFC’s abilities to encode, store, and access multiple schemas. The task requires a recognition of “rule scenarios” (a form of “set learning”) and flexible adaptation through reinforcement signals to shift with changing rules. Patients with prefrontal damage often have difficulty with this task, with some stereotypically making perseveration errors, indicating an inability to switch rules when given reinforcement [4],[5].

Although many models are able to solve the classic WCST (Fig 1c–d and Supplementary Fig. 6), we sought to use the WCST to help uncover the mechanisms by which the PFC is able to learn and remember multiple schema in the absence of curated training or supervision. The framework we develop can be generalized to many similar tasks.

### The model: dynamic mixture of experts (DynaMoE)

Our neural network architecture combines RNNs used previously to model the function of PFC [8] with the MoE architecture [18], and introduces two new features that enable flexible lifelong learning of disparate schemas: a new learning process and repeated focal retuning. Our MoE design uses two specialized networks: a gating network that receives input from the external environment and outputs a decision of which expert network to use; and expert networks that take external input and output an action decision—the card stack to sort the current card in the WCST (Fig. 1a). To capture the complex dynamics of the PFC, we modeled both the gating network and expert networks as RNNs (long short term memory networks (LSTMs) in our implementation). While other architectures have been used in MoE networks, recent work by Wang et al. [8] demonstrated the ability of RNNs to reliably store biologically realistic function approximators when trained by reinforcement learning that mimic animal behaviors.

Using this network architecture, we first introduce a new learning process (Fig. 2b). Our neural network begins as a gating network with a single expert network (Fig. 2a). As it gathers experience in the world, it learns in series of 2-step tunings. When the neural network experiences a scenario (e.g. a series of card sorts in the WCST), it first tunes its gating network to attempt to solve the problem by optimally delegating to expert networks, much as a traditional MoE model would. If some combination of expert actions results in satisfactory performance, no further learning is necessary. If, however, the experiences are sufficiently novel such that no combination of the current expert networks’ outputs can solve the task fully, the network then brings online a latent untrained expert (Fig. 2b–c). The new expert is trained along with the gating network, resulting in a new expert that handles those problems that could not be solved with previous experts. Importantly, this training procedure is agnostic to the order of training scenarios presented and does not require any supervision. Instead, given only the desired performance criteria (e.g. level of accuracy) and limit of training duration per step (how long to try to solve with current experts), our neural network dynamically tunes and grows to fit the needs of any scenario it encounters (Fig. 2c). The learning curves in Fig. 2c reveal two prominent features. First, the speed of learning is successively faster for the second (color sorting) and third (number sorting) scenarios, which is a form of transfer learning. Second, after learning all three scenarios, relearning the first scenario (shape sorting) was rapid, a form of memory savings.

**Figure 2.**
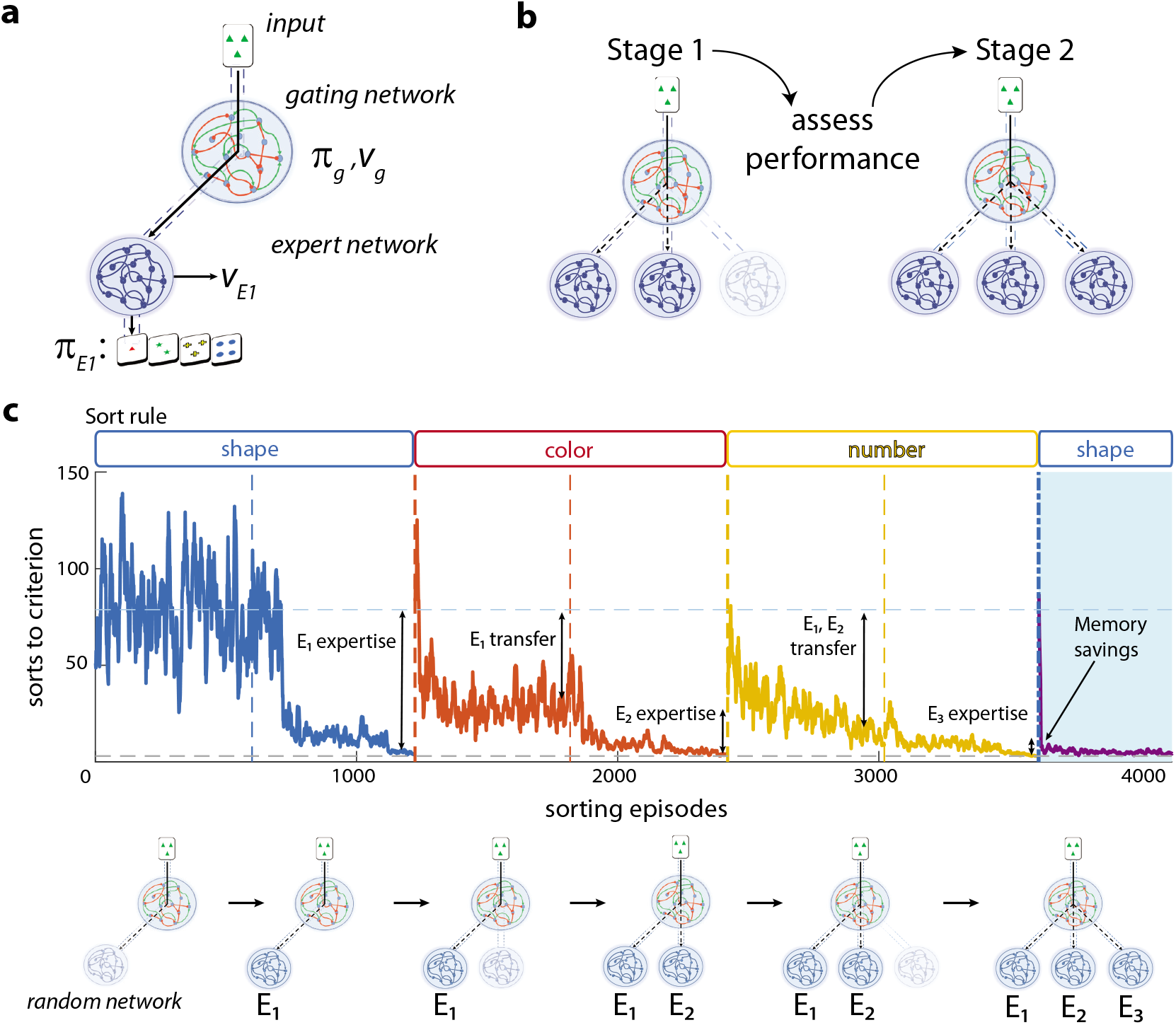
Training of a DynaMoE network. **a**, DynaMoE begins with a gating network and a single expert network, *E*_1_. Both the gating and expert networks train by reinforcement learning, outputing a predicted value (*v_g_* and *v*_*E*1_) and policy (*π_g_* and *π*_*E*1_). **b**, DynaMoE’s 2-step learning process. In Stage 1 the gating network retunes to attempt to solve the task at hand with current experts; if performance is unsatisfactory, the network adds an additional expert in Stage 2 which preferentially trains on tasks that could not be solved by other experts. **c**, A sample training trajectory of a DynaMoE network presented with sequential periods of sorting rules in the WCST. A randomly initialized DynaMoE begins in the shape sorting scenario. First the gating network is tuned alone. In the second step of learning, the first expert network, *E*_1_, is trained (second half of the blue curve). The sort rule is switched to color (red curve) and the same 2-step training process is repeated; followed by the number sort rule (yellow). The improved performance between the first and second stages of training in each sort rule scenario results from expert training. The improved performance from gate retuning results from transfer learning from past experts and increased network capacity. The purple curve shows how DynaMoE rapidly “remembers” past experience due to robust memory savings. The schematic below the graph shows the progression of the DynaMoE network as it experiences the scenarios. Each stage of training above was done for 625 sorting episodes to display convergent learning behavior.

The second new feature is repeated retuning of the gating network. Training standard neural networks on new tasks leads to overwriting of network parameters resulting in catastrophic forgetting [11] (Supplementary Fig. 1). By decomposing a single network into a hierarchy of gating and expert networks, we are able to separate the memory of the neural network into the “decision strategy” (gate), which maps between inputs and experts, and the “action strategies” (experts), which map from input to actions. The hierarchical separation enables repurposing expertise through combinatorial use of previously acquired experts and a natural means to confine memory overwriting to a small portion of the neural network that can be easily recovered through repeated retuning. This results in memory savings [24] that remain robust to new learning and lead to rapid “remembering” rather than relearning from scratch (compare purple curve in blue shaded region in Fig. 2c and Supplementary Fig. 1).

We found that the implementation of these two new features in a hierarchical MoE composed of RNNs results in an architecture that organically learns by reinforcement relative to past experiences and preserves memory savings of past experiences, reminiscent of prefrontal cortex. Importantly, when presented with the classic interleaved WCST, our network (Fig. 1e) learns just as fast or faster than standard RNNs and traditional MoE networks (Fig. 1c–d and Supplementary Fig. 6). We next sought to understand how our dynamic architecture enabled the observed transfer and savings.

### Transfer learning: DynaMoE seeded with pretrained experts

To probe how the DynaMoE network implements transfer learning, we first created an easily interpretable scenario in which two expert networks were separately pretrained on specific rule sets of the WCST; one on shape sorting (*E_shape_*) and another on color sorting (*E_color_*) (Fig. 3a). We then seeded a DynaMoE network with the two pretrained experts and a randomly initialized untrained expert, and introduced it to the third rule, number sorting, and studied its behavior.

**Figure 3.**
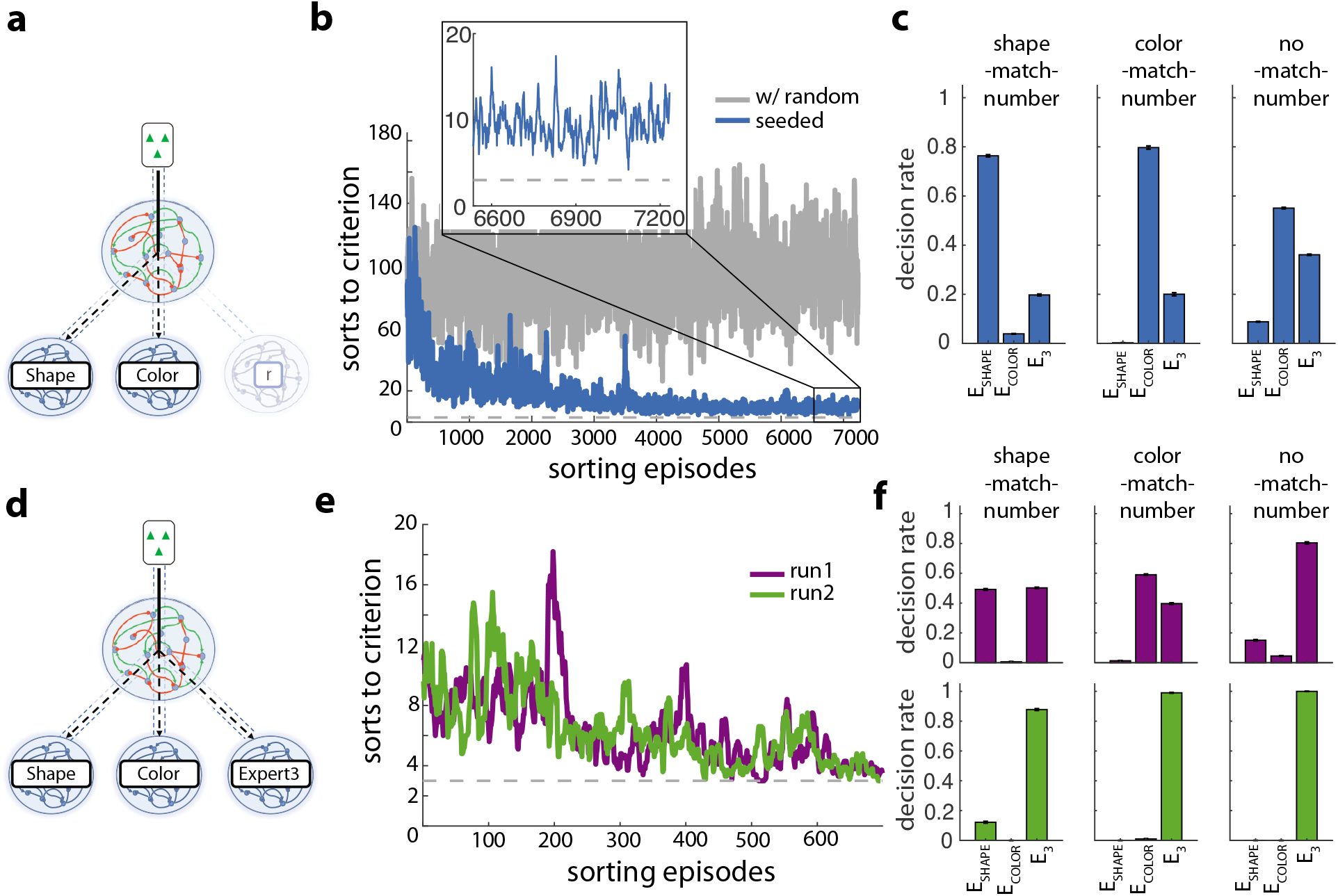
Transfer learning with a seeded DynaMoE network. **a**, A DynaMoE network seeded with pretrained shape and color experts and a randomly initialized untrained network. **b**, The DynaMoE network from **a** achieves near perfect performance in number sorting when only the gating network is trained (blue) in contrast to a network with only an untrained expert (grey). Inset shows that performance of the seeded network does not reach the minimum sorts to criterion (grey dash) without training the third expert network. **c**, The proportion of cards allocated to each expert network after the training in **b** in three different subsets of the number sort rule: shape-match-number, color-match-number, and no-match-number. **d**, Seeded DynaMoE network with trained Expert3 network. **e**, Performance (measured by decrease in sorts to criterion) of two example training runs from the same initial network (**a–c**) that result in different end behavior (**f**). The performance of both runs improves from DynaMoE with an untrained expert (**b** inset) and are indistinguishable from each other. **f**, Top, proportion of experts used in same subsets of number sort rule as in **c** for an example run (run 1). A varying decision rate for experts is used depending on the scenario subset. Bottom, same as above but for a second example run (run 2). The new expert (*E*_3_) is used regardless of subset of number sort rule. See Supplementary Fig. 2 for all 10 runs. Error bars are standard deviation over 1,000 test episodes after training. Absence of bar indicates zero selections of the given expert during testing.

Reflexively, one may speculate that a DynaMoE network with a shape sorting expert, a color sorting expert, and a random network would perform no better on number sorting than with only a random network, since number sorting is seemingly independent of shape or color. Somewhat surprisingly, we found this was not the case. After tuning the gating network, the DynaMoE network with pretrained experts performed drastically better than without them, nearly reaching perfect performance (Fig. 3b). We found that the gating network learned to identify cards for which the shape or color sort matched the correct number sort, and allocate them to the corresponding expert. For example, a card with 1 blue triangle would be sorted to Stack 1 in both the shape (triangle) and number (one) scenarios (“shape-match-number”). Similarly, some cards, for example the card with 1 red circle, would be sorted to Stack 1 in both the color (red) and number (one) scenarios (“color-match-number”). The gating network learned to map these cards to *E_shape_* and *E_color_* to perform correct card sorts in the number rule (Fig. 3c). Only cards for which the number sort did not match the shape or color sort (“no-match-number”) were unsolvable with either *E_shape_* or *E_color_—*for these cards, the gating network used a mixture of the shape, color, and untrained expert networks (Fig. 3c rightmost panel), since no expert could reliably sort these cards correctly. The network had learned to exploit a hidden intrinsic symmetry between the features in the task to enhance performance.

Consequently, when the new expert network was brought online and trained (Fig. 3d), the gating network allocated a large proportion of “no-match-number” cards to the new expert (*E*_3_) (Fig. 3f, rightmost panels). *E*_3_’s expertise thus became number sorting cards that do not match shape or color sorts. Interestingly, this demonstrates a form of transfer learning. The gating network learned to use existent experts to find partial solutions for new problems, leaving unsolvable parts of the problem to be disproportionately allocated to the new expert in the second step of training. New experts thus learn relative to old experts, building on prior knowledge and experience.

In practice, the expertise of *E*_3_ varied between number sorting predominantly “no-match-number” cards and all cards. This likely reflects a trade-off between the complexity of mapping functions the gating and expert networks must learn. In the number sorting scenario, the gating network can learn to map each card type to the appropriate expert or the simpler function of mapping all cards to *E*_3_; *E*_3_ in turn learns to number sort only “no-match-number” cards or all cards. This highlights a trade-off that occurs in biological systems like the brain. We may be able to solve a new problem by piecing together numerous tiny bits of completely disparate strategies, but as complexity of the mapping function increases, at some point it becomes more efficient to simply learn a separate strategy for the new problem, allocating dedicated memory for it.

We found that in the first stage of training, tuning of the gating network consistently led to a mapping function that allocated the vast majority of “shape-match-number” cards to *E_shape_* and “color-match-number” cards to *E_color_* (Fig. 3c). “No-match-number” cards were allocated between all three experts. After the second stage of training in which both the gating network and E3 are trained, we found that the “no-match-number” cards were almost entirely allocated to E3 as expected (Fig. 3f rightmost panels). We found that usage of experts for “shape-match-number” and “color-match-number” cards varied across different training runs (Fig. 3f and Supplementary Fig. 2). To see how often training led to different expert network decision rates, we ran the second stage of training 10 times from the same initial network that had gone through the first stage of training. Usage of the relevant pretrained expert (e.g. *E_shape_* for “shape-match-number” cards) ranged from as much as 65% to as low as 1%, representing end behavior in which *E_shape_* and *E_color_* continued to be used or in which E_3_ was used almost exclusively (run1 and run2 in Fig. 3f, respectively). The non-relevant expert (e.g. *E_color_* for “shape-match-number” cards) was rarely ever used (0–5%). This shows that while DynaMoE networks support pure transfer, the degree of transfer learning implemented depends on network capacity, learning efficiency, and the stochastic nature of learning. All networks achieved the same near perfect performance stop criteria within similar numbers of sorting episodes (Fig. 3e; see Methods).

### Transfer learning: organic case

To probe how a DynaMoE network naturally implements the transfer learning described in the previous section, we trained 10 DynaMoE networks independently from scratch through sequential experiences of the different rules of the WCST. Each network began with an untrained gating network and expert network (*E*_1_). The DynaMoE networks were then trained on shape followed by color and then number sorting, adding a new expert in each new sort rule scenario (see Methods).

As expected from the result in the previous section, we found that the expert networks were not pure rule sorters, but rather had learned an expertise in a mixture of rule types relative to the other experts. For each sort rule scenario, one expert network was used preferentially (*E*_1_ for the first rule experienced, *E*_2_ for the second, etc.), which we refer to as the “dominant expert network” for that sort rule scenario. To quantify the degree of transfer learning utilized, we measured the usage of all 3 expert networks in the different sorting scenarios. For each sort rule scenario, the gating network was retuned until “expert performance” was once again attained. We then measured the usage of each of the non-dominant expert networks with respect to usage of the dominant expert network. Although the magnitude of relative usage varied between independent runs, a consistent pattern emerged. In the shape sort scenario—the first scenario encountered with only *E*_1_—*E*_2_ and *E*_3_ were used very little or never (Fig. 4a, left). For the second scenario encountered—color sort scenario—*E*_1_ was used a small amount, and *E*_3_ was never used (Fig. 4a, middle). Finally, for the third scenario—number sort scenario—*E*_1_ and *E*_2_ were used a small but significant portion of the time (Fig. 4a, right).

**Fig. 4.**
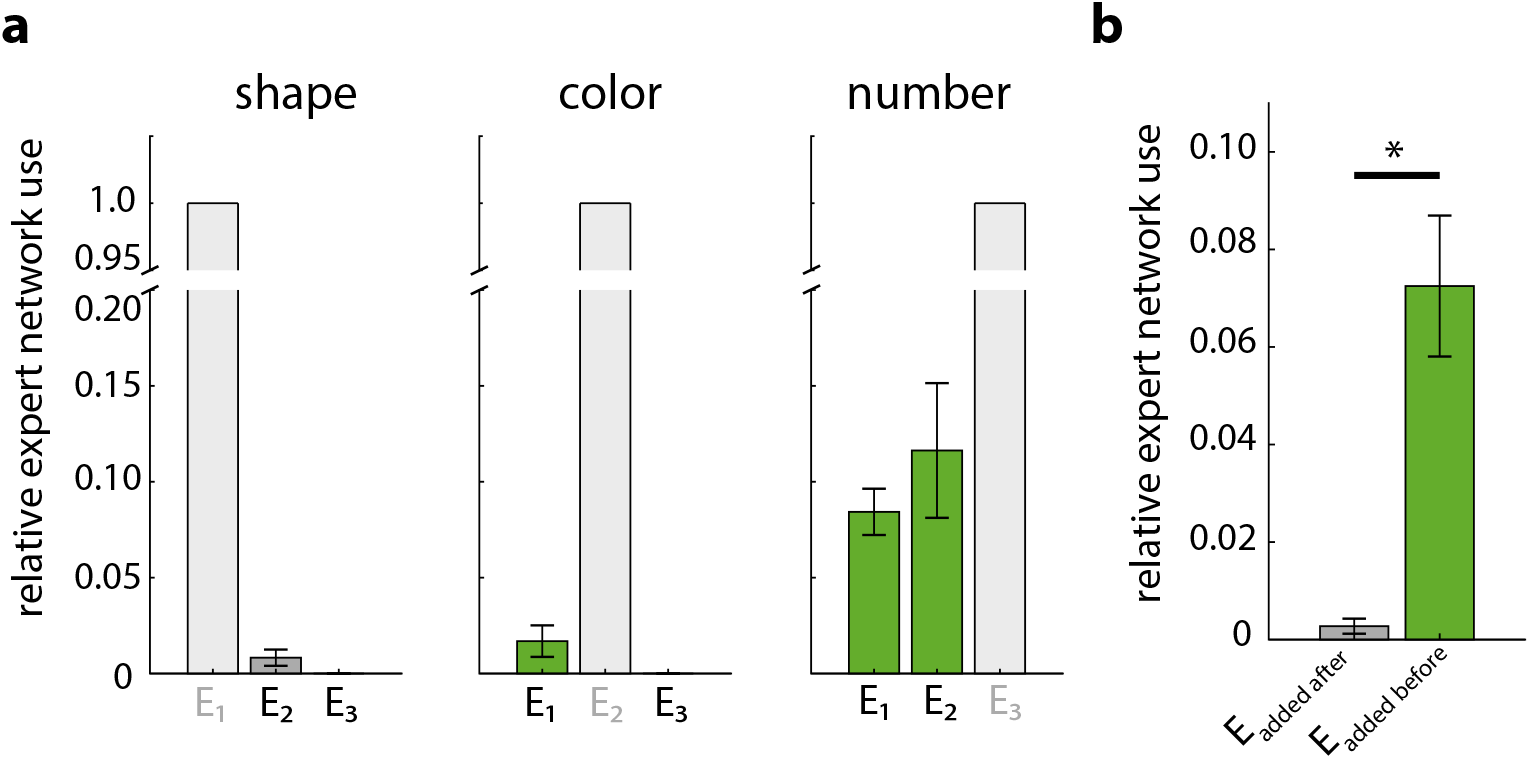
Transfer learning in an unseeded DynaMoE network. **a**, The relative use of each expert network in each sort rule normalized to the dominant expert for the sort rule from 10 independent DynaMoE networks trained in a sequential training regimen (see Methods). The greyed out expert network label with lightest grey bar of value 1 indicates the dominant expert network for each sort rule. Darker grey bars indicate usage of experts that were not present during initial training of the given sort rule (e.g. *E*_2_ and *E*_3_ for the shape rule). Green bars indicate experts that were present during initial training of the given sort rule (e.g. *E*_1_ and *E*_2_ for the number rule). Absence of a bar indicates the given expert was never used. Error bar is SEM over 10 independent runs (see Supplementary Fig. 3). **b**, Aggregated bar plot from **a** grouped by whether the expert was added before or after initial training on the rule. Use of experts present during initial training of a rule indicates transfer learning (green bar), while use of experts not present during initial training indicates non-transfer usage (grey bar). The usage of experts added before was significantly higher (*p* = 1.16*e* – 05, Student’s *t*-test) than that of experts added after initial training on a rule. Error bar is SEM.

This trend of increased usage of experts that were present during the learning of a rule compared to experts added afterward strongly indicates transfer learning as the DynaMoE network encountered new scenarios (Fig. 4b). Newly added experts predominantly trained on examples that the other experts could not solve. Thus when the gating network was retuned to solve a scenario later, it continued to use the previously added experts. In contrast, if an expert was added after the learning of a scenario, all the knowledge to solve the scenario was already contained in the existent experts, so the expert added after learning was rarely used. This shows that new experts were trained relative to knowledge contained by existent experts. Furthermore, while the aggregated expert use percentages clearly show the presence of transfer learning, they mute the degree of transfer learning adopted by some individual networks (Supplementary Fig. 3).

### Robust memory savings

A critical feature of the PFC is the ability to retain knowledge from past experiences. Many neural network models suffer from castastrophic forgetting [11], overwriting information from previous experiences. Put in terms of network parameters, when such networks retune weights to solve new problems, they move to an optimal point in weight space for the current task which can be far away from the optimal space for previous tasks.

In contrast, DynaMoE networks, like the PFC, maintain near optimal configuration for previously experienced scenarios, exhibiting “memory savings” [24]. The hierarchical architecture of DynaMoE networks confines memory loss to a small flexible portion of the network: the gating network. If a scenario has been encountered before, retuning the gating network to optimal configuration is rapid, requiring only a small number of reinforcement episodes (Fig. 5a–b and Supplementary Fig. 5). Retuning the gating network requires much less movement in weight space compared to standard RNNs, since tuning is confined to only the gating network. This is in stark contrast to standard neural networks which can require complete retraining (Fig. 5c–d and Supplementary Fig. 4).

**Fig. 5.**
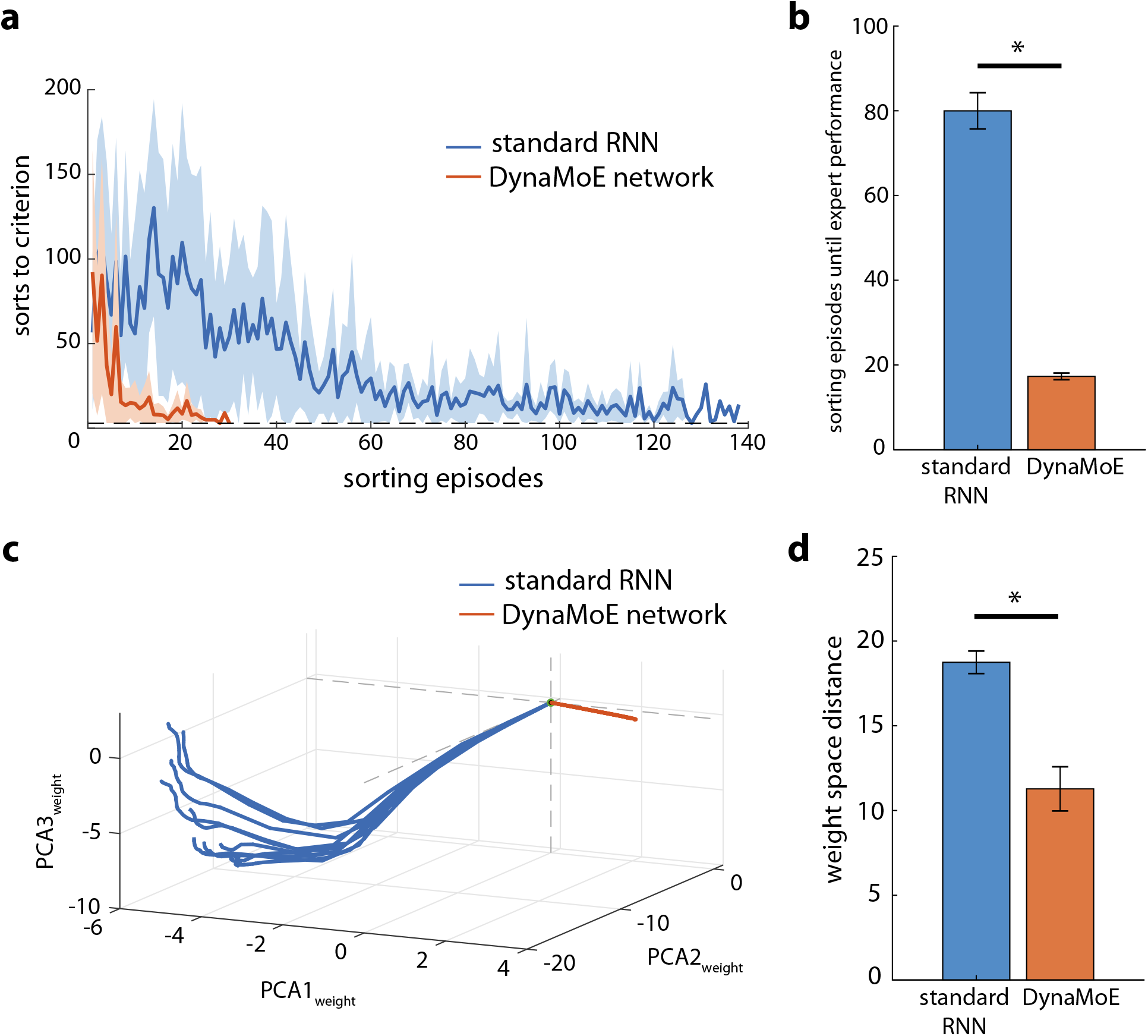
Robust memory savings of DynaMoE. **a**, Example of performance over sorting episodes of retraining of standard RNN (blue) and DynaMoE networks (orange) on a previously encountered task. Shading indicates standard deviation over 10 independent retraining runs of a sequentially trained network. **b**, Average number of sorting episodes required until expert performance for standard RNN and DynaMoE networks over 10 independently trained networks of each type. DynaMoE networks require 78% fewer sorting episodes to remember (*p* = 2.49e —11, Student’s *t*-test). **c**, Visualization of top three principal components of weight space for 10 relearning/remembering trajectories of an example standard RNN and a DynaMoE network trained sequentially. **d**, Euclidean distance between networks before and after remembering previously learned rule in full weight space (average of 10 independently trained networks of each type). DynaMoE network moves 40% less in weight space compared to the standard RNN (*p* = 7.43*e* — 05, Student’s *t*-test). Error bars are SEM.

To measure the memory savings of DynaMoE networks, we sequentially trained networks with identical presentations of, first shape, then color, then number sorting scenarios (see Methods). We then tested how many sorting episodes of reinforcement were required for the network to regain expertise in the first sorting rule it experienced (shape). As Fig. 5a–b shows, DynaMoE networks required 78% fewer episodes to regain expertise than standard RNNs (*p* = 2.49*e* — 11, Student’s *t*-test). The number of episodes required to remember was drastically fewer than when it first learned the rule, whereas standard RNNs improved only slightly compared to when they first learned the rule (Supplementary Figs. 1, 5, and 6). While standard RNNs nearly completely overwrote the information learned through initial training, DynaMoE networks preserved their memory and only required brief reinforcement for the gating network to remember how to allocate cards to experts.

To measure the weight changes required to regain optimal performance, we measured the distance in weight space each of the networks traversed when remembering the shape sort rule after sequential training. DynaMoE networks traversed 40% less distance in weight space to reach optimal performance compared to standard RNNs (*p* = 7.43*e* — 05, Student’s *t*-test; Fig. 5c–d and Supplementary Fig. 4). Even after sequential training, DynaMoE networks remain relatively close in weight space to the optimal performance configurations on all previously experienced tasks. In contrast, standard RNNs moved far from their initial optimal point in weight space for the shape scenario, resulting in movement of nearly equal distance when relearning the shape scenario as when initially learned (Supplementary Fig. 4).

### Lesions of DynaMoE cause PFC lesion-like impairments

The DynaMoE framework provides an opportunity to understand how disruptions to specific functional aspects of the PFC and related areas can lead to different impairments observed in clinical cases. Numerous clinical and neuroimaging studies have indicated regional specialization within the PFC, yet evidence from human studies is invariably messy, involving overlapping brain regions and varying degrees of impairment in different aspects of tasks [2],[5],[25]. Our framework enables targeted disruption of specific functional components of our network that may help clarify the underlying organization of the human PFC. The WCST has served as a standard clinical assessment to evaluate PFC impairment [5] making it an ideal task with which to analyze functional consequences of various lesion types.

To assess how lesions of our network architecture could result in behavioral impairments, we damaged specific regions of the gating network of our architecture. Importantly, in our lesion studies the expert networks were unperturbed, leaving available the action strategies to perfectly perform the task. This characteristic is often seen in patients with prefrontal damage: although they have difficulty with the full WCST, if explicitly told which sort rule to use, patients are often fully capable [5],[26]. We first trained DynaMoE networks on each rule type, and then the classic interleaved WCST (Supplementary Fig. 6e; see Methods). We then lesioned the gating network and performed testing on the classic WCST to assess changes in performance and behavior.

Lesions were targeted to five different regions within the gating network (Fig. 6a): inputs to the network (red region 1 in Fig 6a)—ablation of reward feedback (L1), action feedback (L2), or both (L3); internal network dynamics (region 2)—ablation of varying numbers of synaptic connections to the “forget” gate of the LSTM (L4_0.5_ and L4_0.9_ were 50% and 90% ablations of synaptic connections, respectively; Supplementary Fig. 7a–b for full range); output of the network (region 3; L5); and areas downstream of the network—ablation of synaptic connections to the units that determined which expert network to use (region 4; L6), to the unit that estimated value (region 5; L8), or both (L7; Supplementary Fig. 7c–d for full range). Since lesions could potentially have differential effects depending on the specific disruptions incurred, we performed each lesion 10 times and measured the average effect.

**Fig. 6.**
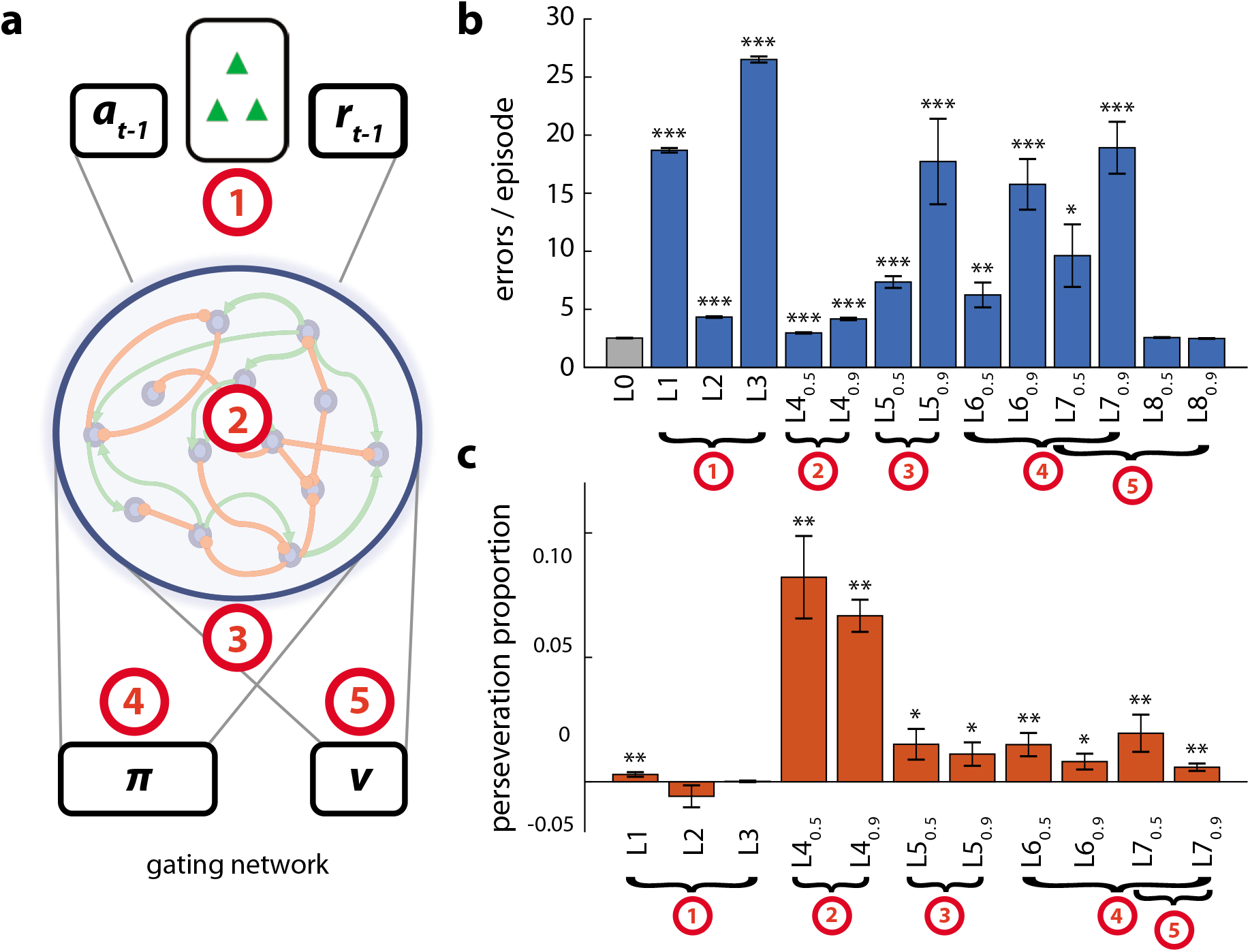
Different lesion-induced error modes of DynaMoE gating networks. **a**, Map of lesioned regions in DynaMoE’s gating network. Three lesions of input (region 1) were done (L1–L3), one lesion of the network dynamics (region 2—L4), one lesion of network output (region 3—L5), one lesion of decision determination (region 4—L6–L7), one lesion of value determination (region 5—L7–L8). **b**, Average number of errors per episode for each lesion type. L1 is ablation of reward feedback from previous trial; L2 is ablation of action feedback from previous trial, L3 is simultaneous L1 and L2 lesions; L4_0.5_ is ablation of 50% of connections to forget gate of network and L4_0.9_ is ablation of 90%; L5 is ablation of output units; L6 is ablation of connections to decision units (π); L8 is ablation of connections to value unit (v); L7 is simultaneous L6 and L8. Asterix indicates significant difference from no lesion (L0) (Student’s *t*-test *: *p* < 0.05; *: *p* < 0.01; ***: *p* < 0.001) **c**, Proportion of increase in errors that were perseveration errors for lesions that caused significant increase in errors. Asterix indicates confidence interval (CI) excluding zero (*: 95% CI; **: 99% CI). All error bars are SEM.

We found that different lesion types resulted in different degrees of impairment, ranging from no change in error rate (e.g. L8_0.9_, *p* = 0.3664, Student’s *t*-test) to 10.5 fold more errors (L3, *p* = 2.1268*e* — 25, Student’s *t*-test) than before the lesion (Fig. 6b and Supplementary Table 1). Since perseverative errors are a signature of some prefrontal lesions, particularly associated with dorsolateral PFC (dlPFC) impairment, we measured the proportion of the increase in error rate that was due to perseverative errors (“perseveration proportion”). Fig. 6c shows the variability in perseveration proportion for the lesions that caused a significant increase in error rate.

The impairment profiles caused by lesions to specific functional components characterized by increase in total error and perseveration proportion reveal different error modes that mirror the different error modes observed from patients across the range of prefrontal lesions (Fig. 6 and Supplementary Table 1). Overall, lesions grouped qualitatively into three categories: lesions that caused often substantially increased total error rate (1.72-10.51x), a small proportion of which were perseverative errors (0-1.95%) (regions 1,3,4; L1-L3,L5-L7); lesions that caused a small but significant increase in total errors (1.18-1.65x), a large proportion of which were perserverative errors (6.67-8.22%) (region 2; L4); and lesions that caused no change in error rate (region 5; L8).

Our lesion results provide a roadmap with which to interpret and understand the variety of error modes observed in human patients with prefrontal damage due to trauma or disease. While the PFC as a whole has been definitively linked to set-shifting and cognitive flexibility, localization of functional components to specific subregions remains unclear. Lesions throughout the prefrontal areas have been associated with impairments observed in the WCST, ranging from no change in error rate to large increases in perseverative and non-perseverative error rates similar to the range of behavioral outcomes resulting from our lesions [5]. Canonically, though with mixed evidence, impairment of the dlPFC is associated with increased error rate on the WCST, particularly perseveration errors. Our lesion study indicates this behavioral phenotype maybe due to impairment of gating network dynamics, suggesting the dlPFC may contribute to a gating-like mechanism within the PFC.

## Discussion

In this paper we propose a new framework for how the PFC may encode, store, and access multiple schemas from experiences in the world. Like the PFC, the DynaMoE neural network is agnostic to training regimen and does not require “oracle” supervision. We showed how the hierarchical architecture of DynaMoE naturally leads to progressive learning, building on past knowledge. We then demonstrated how DynaMoE networks reliably store memory savings for past experiences, requiring only brief gate retuning to remember. Finally, we showed how lesions to specific functional components of the DynaMoE network result in different error modes in the WCST, analogous to the error modes described of patients with different forms and severity of prefrontal damage.

The parallels seen between the DynaMoE network and the PFC and related areas encourages investigation into the extent to which these two systems recapitulate each other. Perhaps most poignantly, this comparison puts forth the hypothesis that the PFC may be organized as a gating system that is tuned to optimally combine knowledge from past experiences to handle problems as they are encountered. Some studies have provided evidence for such a functional architecture in the brain [27], and prefrontal cortical areas in particular [28],[29],[30]. Experimental investigations that compare the neural activity in DynaMoE networks to that in prefrontal cortical areas will be fruitful in supporting or refuting this hypothesis and is a topic we are currently exploring.

Our lesion studies motivate further investigation of functional specialization in the PFC through comparison of our framework and clinical, experimental, and neuroimaging studies [2],[5]. Clinical and experimental studies have yielded unclear and sometimes contradictory findings due to the anatomical inseparability of PFC functions [5]. Our model provides full access to the underlying structure enabling targeted studies to use as a reference for interpreting humans studies. Further comparison of our framework with in-depth phenotypic analyses across various tasks may help us understand the functional organization of the PFC and the consequences of disruptions due to trauma and disease.

Our lesion analysis also motivates future studies on adaptation to lesions. In the present study we focused on lesions after learning was complete, since most clinical case reports describe testing of patients after acute injury. In clinic, it is also important to understand how patients may cope and adapt after a lesion has occurred. The DynaMoE framework may be useful for studying the effects lesions on learning and adaptation.

The DynaMoE framework also has interesting implications for areas of machine learning. Its organic, unsupervised implementation of transfer may be useful for intractable problems that may be handled piece-wise in a way that may be non-obvious to an “oracle” supervisor. By letting the model learn how to grow and structure itself, our framework puts the burden of optimally solving complex problems on the algorithm. This may significantly improve progress by removing the need for careful curation of training data and training regimen.

The form of transfer learning demonstrated by our dynamic architecture—acquiring new knowledge (new expert) based on indirect knowledge of what other parts of the network (old experts) know—has not been reported before to our knowledge. This form of transfer learning is reminiscent of “learning by analogy,” a learning skill humans are very good at, but machines continue to struggle with [31],[32]. Through our framework, this dynamic form of transfer could be extended to much larger networks, utilizing a myriad of experts. Such a framework could be useful both as a model of the brain and for machine learning applications.

Finally, our framework provides a new method for lifelong learning and memory. Major challenges persist in developing methods that do not get overloaded, but also scale well to lifelong problems [33]. Similar to “grow-when-required” algorithms, our network adds capacity when necessary. However, our network also leverages already acquired knowledge to help solve new problems, reducing demand for growth. This feature supports scalability, which both the brain and machine learning methods must support given their limited resources. Elaborating and adapting DynaMoE to more complex tasks and incorporating other techniques such has simultaneous combinatorial use of experts will lead to exciting steps forward in lifelong learning.

## Methods

### Behavioral task

To demonstrate our framework we used the Wisconsin Card Sorting Task. In this task the subject is asked to sort cards with symbols. Each card has symbols with a shape type (triangle, star, cross, circle), a color type (red, green, yellow, blue), and a specific number of symbols (one, two, three, four). During each episode, an unsignaled operating rule is chosen: either shape, color, or number. The subject must discover the rule by trial and error and then sort a given card according to the relevant rule into one of four stacks. The first stack, Stack 1, has 1 red triangle, Stack 2 has 2 green stars, Stack 3 has 3 yellow crosses, and Stack 4 has 4 blue circles. For example, a subject could be given a sample card with 3 green triangles. If the operating rule is shape, the card should be placed in Stack 1 (matching the triangle shape of the 1 red triangle). If the operating rule is color, the card should be placed in Stack 2 (matching the green color of the 2 green stars). If the operating rule is number, the card should be placed in Stack 3 (matching the number of the 3 yellow crosses)(Fig. 1b). After each attempted card sort, the subject is given feedback as to whether the sort was correct or incorrect. Once the subject has sorted a given number of cards correctly consecutively the operating rule is switched without signal and the subject must discover the new rule through trial and error. For all of our simulations, the operating rule was switched after 3 correct sorts in a row.

At the beginning of each episode a deck of cards containing all 64 possible combinations of shape, color, and number was generated. Cards were randomly drawn from this deck and presented to the subject for sorting, removing each card from the deck after presentation. If all 64 cards from the deck were used before termination of the episode, the deck was regenerated and new cards continued to be drawn in the same manner. An episode was terminated by meeting one of two termination criteria: 1) achieving the given number of correct sorts in a row (3 for our simulations) or 2) reaching the maximum episode length which we set to 200 card draws.

In our sequential scenario training simulations, a particular operating rule was kept constant for the duration of training in that period, either until a given number of sorting episodes was achieved or until performance passed a satisfactory threshold. In the next training period, a new operating rule was held constant and training was repeated in the same manner. As a demonstration, a DynaMoE network was trained in a sequential training protocol with sequential blocks of 1,250 sorting episodes (1 “sorting episode” ≈ 12 total WCST episodes across whole network) of each sort rule type (Fig. 2c). Each 1,250 sorting episode block was split into two 625 sorting episode subblocks; in the first subblock, the gating network was tuned and in the second both the gating and new expert network were tuned. When the shape sort rule was reintroduced, only the gating network was tuned. The 1,250 sorting episode block training protocol described above was also done with a standard RNN for comparison (Supplementary Fig. 1). In all line plots of sorts to criterion over training, a moving mean over every 10 sorting episodes was calculated and plotted for readability.

### Reinforcement learning training

To train our networks with reinforcement learning, we used the Advantage Actor-Critic algorithm of Mnih et al. [34], where a full description of the algorithm can be found. Briefly, the objective function for our neural network consists of the gradient of a policy term, an advantage value term, and an entropy regularization term:

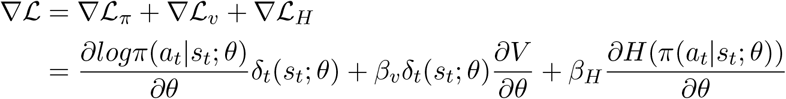

where *π* is the policy, *a_t_* is the action taken at time *t, s_t_* is the state at time *t, θ* is the network parameters, *β_v_,β_H_* are hyperparameters for scaling the contribution of the value and entropy terms respectively, *V* is the value output of the network, and *H* is the entropy regularization term of the policy. *δ_t_* is the advantage estimate, which represents the temporal difference error:

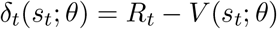

where *R_t_* is the discounted reward:

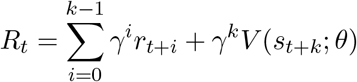

where *k* is the number of steps until the next end state. When *γ* = 0, *R_t_* = *r_t_*.

The advantage equation in this case is equivalent to a temporal-difference error signal enabling temporal difference reinforcement learning.

The parameters of the model were updated during training by gradient descent and back propagation through time after the completion of every 3 episodes. For all simulations we used 12 asynchronous threads for training. In our plots, a single “sorting episode” was defined as the number of total WCST episodes completed while a single thread completed 1 episode, which was roughly equal to 12 episodes for the total network. We used the AdamOptimizer with a learning rate of 1*e*-3 to optimize weights. The objective function scaling hyperparameters *β_v_* and *β_H_* were both set to 0.05 for all our simulations.

For feedback as to whether each card sort was correct or incorrect, we gave a reward of +5 if correct and −5 if incorrect. For the WCST, a discount factor of *γ* = 0 was used since each card sort was an independent event, based only on the relevant operating rule rather than any prior previous action sequence.

Similar to the implementation by Wang et al. [8], the input to the networks for each step was given as vector with the current card shape, color, and number, the action taken for the previous time step, *a*_*t*-1_, and the reward given for previous card-sort action, *r*_*t*-1_.

### Network architecture

Both our standard RNN and DynaMoE network architectures were composed of LSTMs as implemented by Wang et al. [8]. In contrast to “vanilla” RNNs, LSTMs copy their state from each time step to the next by default and utilize a combination of built-in gates to forget, input new information, and output from the states. This RNN structure allows for robust learning and storage of functional approximators for various tasks as demonstrated by Wang et al. [8]. The LSTM states and gates are described by the following equations:

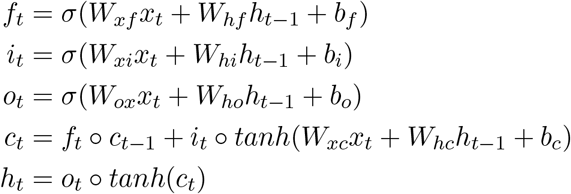

where *f_t_*, *i_t_*, and *o_t_* are the forget, input, and output gates at time *t* respectively, *σ* is the sigmoid activation function, *W_ij_* denotes the weights from component *i* to component *j*, *x_t_* is the external input at time *t, h_t_* is the ouput of the LSTM at time *t, c_t_* is the state of the LSTM cell at time *t, b_f_, b_i_*, and *b_o_* are the biases of the forget, input and output gates respectively, *b_c_* is the bias of the cell states, and o denotes the Hadamard product.

Our “standard RNN” architecture consists of a single LSTM network. As input, the RNN takes the card from the current time step and the action and reward from previous time step. The RNN sends output to a policy layer and a value layer. The policy layer implements a softmax transformation over the possible actions to determine the action to take. The value layer outputs a single number value estimate used to calculate the advantage term for the objective function.

In our DynaMoE architecture, the gating network consists of a single LSTM network. The gating network takes as input the card from the current time step and the action and reward from the previous time step. It sends output to two parallel layers, one implementing a softmax transformation over the possible expert networks to determine which expert network to use, and the other estimating the value of action to be taken. Each expert network consists of a single LSTM network that takes the same input as the gating network, and sends output to two parallel layers. One of these layers implements a softmax transformation over all possible actions to determine the action to take, and the other gives a value estimate of that action. The gating network and the expert networks are each optimized by their own objective function in the form described in the Reinforcement learning training section above.

For all our simulations described in the paper we used a standard RNN of 105 units and a DynaMoE network with a 98 unit gating network and 19 unit experts. We chose these network sizes because they provided ample capacity to learn the WCST scenarios and shared the same number of total trainable network parameters (47,145) which enabled the direct comparisons between standard RNN and DynaMoE networks.

In DynaMoE networks, if the gating network could not solve a scenario using its current experts, a new expert was brought online. In this case, first the gating network was retuned with the current experts and an additional randomly initialized expert of the same size. If performance did not achieve the desired performance criterion, the gating network and the new expert were then trained simultaneously. The gating network LSTM learned a functional approximator mapping from inputs to experts, and the experts learned functional approximators mapping from inputs to actions in their input domain of expertise which was determined by the gating network’s mapping function.

### DynaMoE seeded with pretrained experts transfer simulations

For our demonstration of DynaMoE networks’ transfer learning property, we performed a simulation with pretrained experts. We trained one expert on only shape sorting until the expert network achieved near perfect “expert performance,” defined in this simulation as an average sorts to criterion of <4 in the last 100 episodes of a single asynchronous thread (minimum sorts to criterion is 3). We repeated the same with a second expert network trained on only color sorting. We then created a DynaMoE network with these two pretrained expert networks and a third randomly initialized expert network, and trained the gating network only on the number sorting rule for 7,500 sorting episodes to ensure convergent decision behavior. “Expert performance” as defined above was not achieved during this stage of training (Fig. 3b inset). Network weights were then fixed and behavior and performance of the network was evaluated. To evaluate behavior of the network, 1,000 test episodes were performed in the number rule and the proportion of decisions to use each expert network (the decision rate) was measured in subsets of the number rule described in the the Results section (“shape-match-number,” “color-match-number,”“no-match-number;” Fig. 3c). From this parent network, we then ran 10 independent training runs in parallel in which the gating network and the randomly initialized expert network were trained simultaneously on the number sorting rule until the “expert performance” criteria was achieved. To evaluate the decision rate of the gating network for each of the 10 independent training runs, 1,000 test episodes were performed and the mean and standard deviation of the decision rates were calculated in the same subsets of the number rule (Fig. 3f and Supplementary Fig. 2).

### Organic transfer simulations

For our demonstration of the DynaMoE network’s implementation of transfer learning without any pretraining, we independently trained 10 DynaMoE networks from scratch in the following manner. We began with a randomly initialized gating network with a single randomly initialized expert network. The gating network was then trained alone on the shape sort scenario of the WCST for 1,250 sorting episodes. The gate and single expert network were then trained simultaneously until “expert performance,” defined for this simulation as an average sorts to criterion of <4 in the last 100 episodes of a single asynchronous thread. A new randomly initialized expert network was then added, and the gating network was trained for 7,500 sorting episodes in the color scenario to allow full convergence of decision behavior. The gate and new expert were then trained simultaneously until “expert performance” was achieved. This was repeated finally for the number scenario and a third expert network. To evaluate transfer, for each sort rule we retuned the gating network until expert performance was achieved. The gating network was then tested for 1,000 episodes in the given sort scenario and the relative expert network use was measured as described in the Data analysis section below (Fig. 4 and Supplementary Fig. 3).

### Robust memory savings simulations

To demonstrate the DynaMoE network’s robust memory savings, we independently trained 10 DynaMoE networks and standard RNNs with the same number of trainable parameters (47,145) in an identical presentation of scenarios. First, the randomly initialized networks trained on 1,250 sorting episodes of the shape sort scenario to ensure convergent performance. This was followed by 1,250 sorting episodes of the color sort scenario, followed by 1,250 sorting episodes of the number sort scenario (same as for networks in Fig. 2c and Supplementary Fig. 1). For the DynaMoE network each block of 1,250 sorting episodes with a sort rule was broken into 2 sub-blocks of 625 sorting episodes—in the first 625 sorting episodes, the DynaMoE network did the first stage of training in which only the gating network is tuned, and for the second 625, the second stage of training was done in which both the gating and new expert networks are tuned simultaneously. After this sequential scenario training, for each standard RNN and the DynaMoE network, we ran 10 independent retrainings on the first scenario encountered: the shape scenario. For the DynaMoE network only the gating network was retuned. To measure how quickly the networks could recover performance in the previously learned rule, the networks were tuned until they reached a performance criteria of average sorts to criterion <10 cards for the last 10 episodes of a single asynchronous thread. The number of sorting episodes required to achieve this performance were measured, as well as the distance traveled in weight space during relearning/remembering the shape scenario (Fig. 5 and Supplementary Fig. 4).

To compare how many sorting episodes it took for each network type to relearn/remember the shape scenario after the sequential rule training as it did to learn the scenario for the first time from scratch, we trained 10 randomly initialized standard RNNs and 10 randomly initialized DynaMoE networks with a single expert network until they achieved the performance criteria of average sorts to criterion <10 cards for the last 10 episodes of a single asynchronous thread (Supplementary Fig. 5).

### Classic WCST simulations with untrained and pretrained networks

To simulate performance on the classic WCST in which the different sorting rule episodes are interleaved randomly, 5 different networks were created. The first network was a standard RNN with randomly initialized weights (Supplementary Fig. 6a). The second network was a standard RNN that was pretrained sequentially on first the shape rule, followed by the color rule, followed by the number rule (Supplementary Fig. 6b). For each rule type, the network was trained until “expert performance,” defined as average sorts to criterion <4 over last 100 episodes of single asynchronous thread before switching rules. The third network was a DynaMoE network with three untrained expert networks with randomly initialized weights (Fig. 1c and Supplementary Fig. 6c). The fourth network was a DynaMoE network seeded with three pretrained expert networks—one pretrained on shape sorting, one on color sorting, and one on number sorting (Fig. 1d and Supplementary Fig. 6d). Each of these pretrained experts had been trained on the given rule until reaching “expert performance.” The fifth network was a DynaMoE network that was pretrained sequentially on first the shape rule, followed by the color rule, followed by the number rule (Fig. 1e and Supplementary Fig. 6e). The network started with a gating network and a single expert network with randomly initialized weights. The gating and expert networks were trained simultaneously on the shape rule until “expert performance” was reached. The rule was then switched to the color rule and a new expert network with random weights was added. The gating network was trained for a maximum of 250 sorting episodes and then the new expert was brought online and trained until “expert performance.” The same was then repeated for the number rule.

Each network was then trained on the classic WCST, in which rules are randomly interleaved (rules switch after every episode; see full description in Behavioral task section of Methods). The middle column of Supplementary Fig. 6 shows performance of each network over 2,500 sorting episodes of training. Networks with pretraining (Supplementary Figs. 6b,d,e), were also trained for 2,500 sorting episodes on the shape rule (the first rule experienced) to compare each network’s ability to “remember” a previously learned rule.

### Lesion studies

To perform the lesions studies, we first trained a DynaMoE network identical to the network in Supplementary Fig 6e as described above. We then implemented one of the following lesions: L0- no lesion; L1- ablation of the reward feedback input to the network; L2- ablation of the action feedback input; L3- both L1 and L2 simultaneously; L4- ablation of varying amounts of the synaptic connections of the “forget gate” component of the LSTM, ranging from 10–100% and denoted by the subscript (e.g. L4_0.9_ has 90% of the synaptic connections ablated) (Supplementary Fig. 7); L5- ablation of varying amounts of output from the RNN; L6- ablation of synaptic connections to the units used to determine which expert network to use; L8- ablation of the synaptic connects to the unit used to estimate value; and L7- both L6 and L8 simultaneously. For two of the lesions types (L4,L7) we show the full severity spectrum as an example in Supplementary Fig. 7.

After implementing the lesion, we then tested the full DynaMoE network on the classic interleaved WCST. We ran 1,000 test episodes and then performed analysis on performance as described in the Results section. For each lesion type, we randomly ablated at each level of severity 10 times and analyzed average behavior since lesions of specific connections or units within a given region may have differential effects.

### Data analysis

For the organic transfer learning (without pretraining) analysis, after sequential training, the gating network was retuned for each rule type. The “dominant expert network” was then defined as the expert network that received majority of the decisions after retuning in that scenario. We found the newly added expert was always the “dominant expert network:” first expert for the shape rule, second expert for the color rule, and third expert for the number rule. Of the 10 independent training runs, the minimum usage of the first expert in the shape scenario was 96.45% and maximum of other expert usage was 3.55%; minimum usage of second expert in color scenario was 93.19% and maximum of other expert usage was 6.81%; minimum usage of third expert in number scenario was 67.96% and maximum of other expert usage was 24.34%. To calculate the relative expert network use shown in Fig. 4 and Supplementary Fig. 3 we normalized the non-dominant expert usages to the dominant expert usage in each scenario. The aggregate relative usage of the expert networks in Fig. 4 was calculated by combining the relative usage of all non-dominant experts that were present during initial learning of a rule (first expert for color and number training; second expert for number training) to represent experts present during initial training (green bar), and the relative usage of all non-dominant experts that were not present during initial learning of a rule (second and third experts for the shape training; third expert for the color training) to represent experts not present during initial training (grey bar).

Two analyses were done to assess the networks’ memory savings. In the first, we measured the number of sorting episodes required to achieve expert performance in the first sort rule (shape sort) after going through the sequential rule scenario training described above. We ran this simulation 10 times for both the standard RNN and DynaMoE network, and compared the results with the Student’s t-test. We also used the Student’s *t*-test to compare the number of sorting episodes each network required to relearn/remember the shape scenario to the number of sorting episodes required to learn the shape scenario for the first time (Supplementary Fig. 5).

In the second analysis, we measured the change in weights during retraining of both the standard RNN and the DynaMoE network on the first sort rule (shape) after sequential training. We did this for all 10 repetitions of the retraining. To visualize the movement in weight space for the retraining runs of a network, we performed PCA on the weights for all retraining runs of both networks together. We then took the top three principal components, translated the starting point in this 3 PCA weight space for all networks to the same origin point, and plotted the retraining trajectory of each repetition for each network type (Fig. 5c). For 10 independent full training and retraining runs, we also measured the Euclidean distance between the start and end points in full weight space and compared these using the Student’s *t*-test (Fig. 5d). For the comparison of movement in weight space of the two networks throughout training and remembering, distances traveled in weight space for 10 runs of each type of network were measured throughout the training and remembering periods (Supplementary Fig. 4).

## Acknowledgements

We thank Robert Kim, Yusi Chen, and Gal Mishne for helpful discussions and feedback on the manuscript. We thank Jorge Aldana for support with computing resources. This work was supported by an Office of Naval Research grant no. N000141310672 and National Science Foundation grant no. 1735004.

## Author contributions

B.T. and T.J.S. designed the network architecture and simulation of the behavioral task. B.T. implemented the design, ran the simulations, and carried out the data analysis. K.M.T. and H.T.S. helped frame the manuscript. B.T. and T.J.S. wrote the manuscript.

## Competing interests

The authors declare no competing interests.

## Supplementary Figures/Tables

**Supplementary Figure 1.**
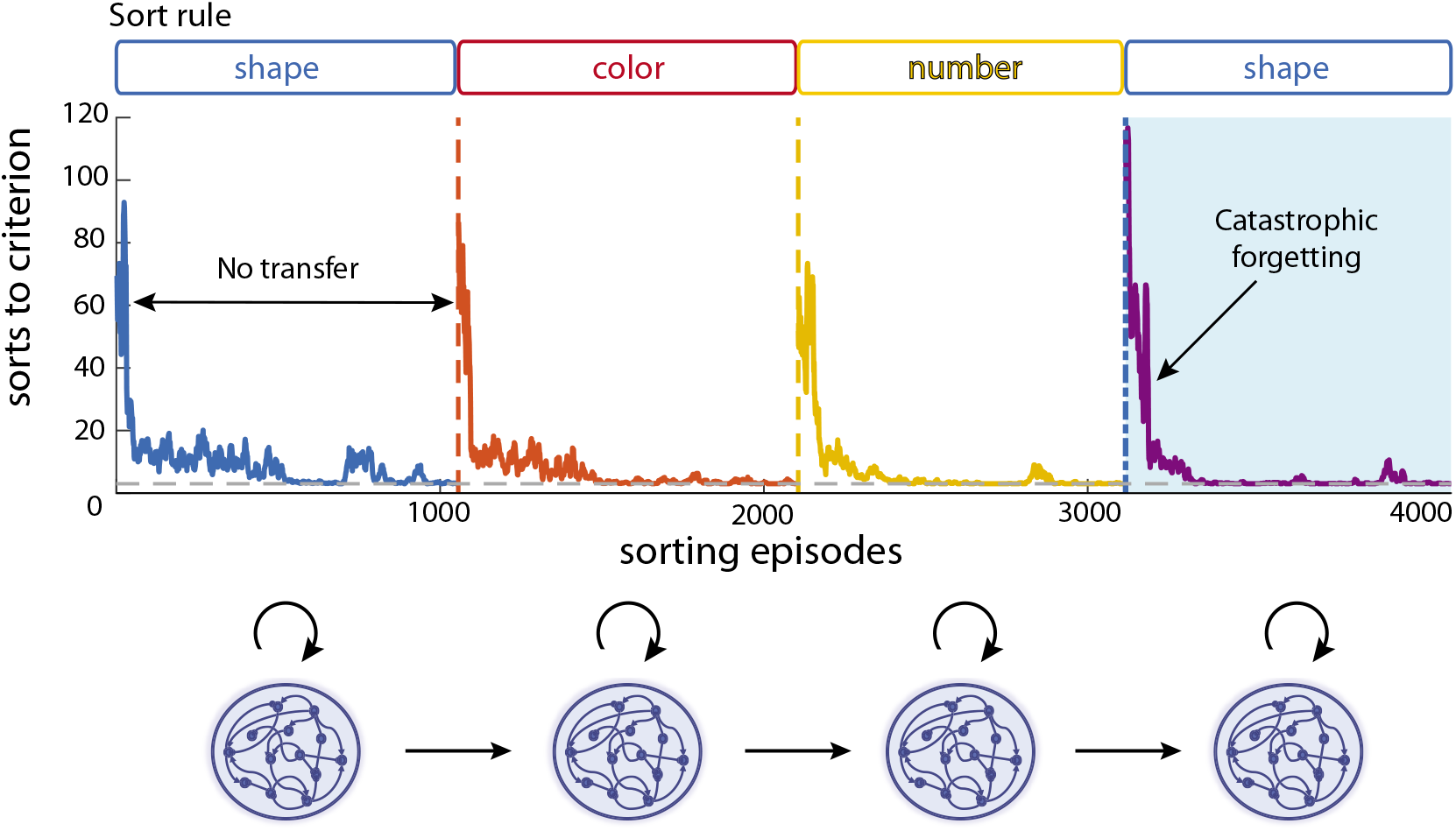
Sequential training of a standard recurrent neural network. For each task, the network was trained for 1,250 sorting episodes to display performance behavior over time (lower sorts to criterion is better performance in the WCST). During learning, all network parameters are tuned (bottom schematic). When trained on a sequences of tasks, the performance trajectory of the network appears nearly identical for every new task, demonstrating the lack of transfer learning for solving new tasks. The nearly identical performance trajectory for relearning a previously learned rule—the shape rule shown in the leftmost white panel and the rightmost light blue—demonstrates the memorylessness of standard RNNs, also called catastrophic forgetting.

**Supplementary Figure 2.**
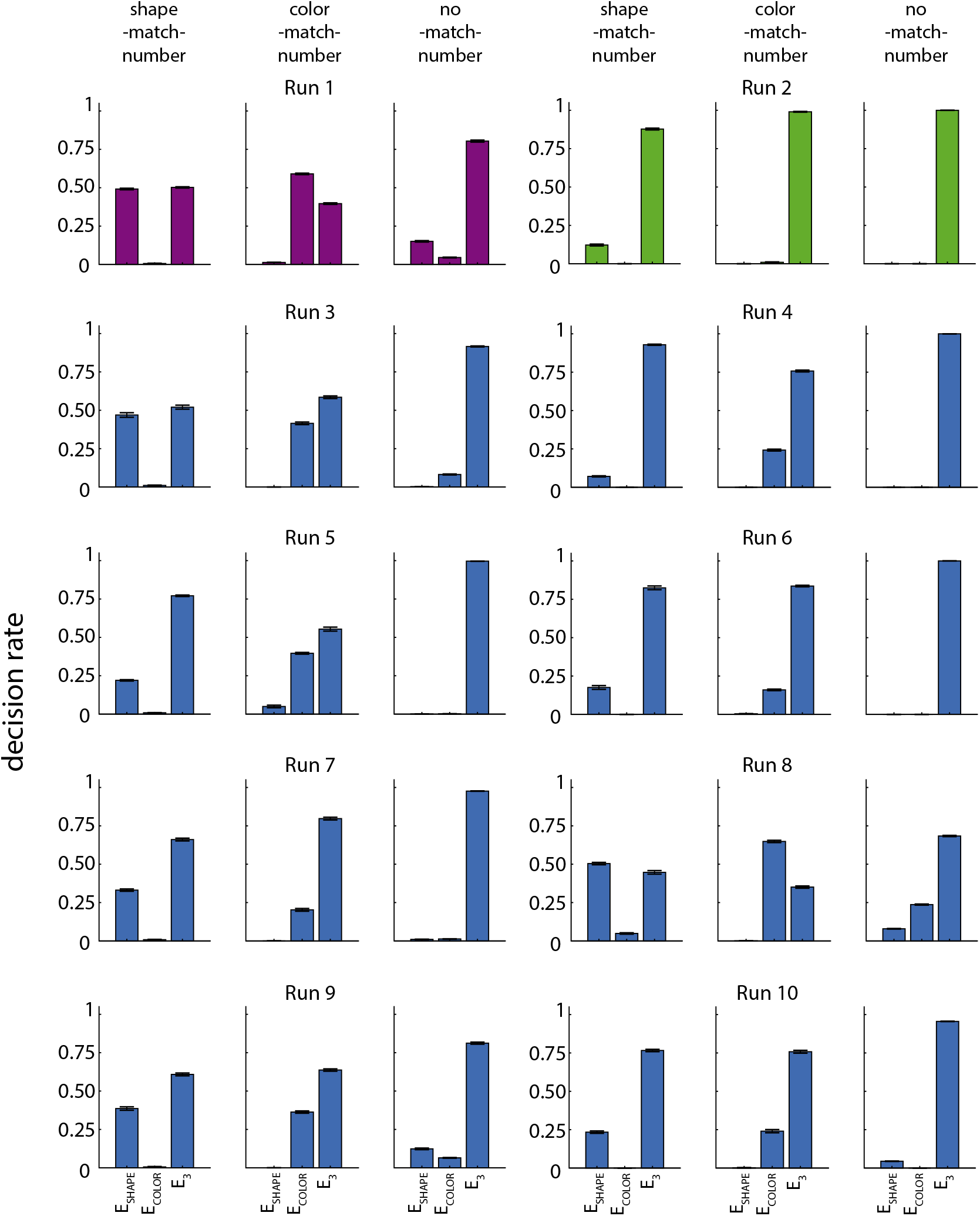
Decision rate in subsets of number sort rule after 10 independent training runs from the same seeded DynaMoE network. For each training run, the gating network and a new expert network (*E*_3_) were simultaneously trained on the number sort rule until “expert performance” (see Methods). Weights were then fixed and 1,000 test sorts were done in the number sort rule. The proportion of decisions by the gating network to use each expert in subsets of the number rule were measured. The purple and green colored runs are the same as depicted in Fig. 3e-f. Error bars show standard deviation over 1,000 test trials. Different training runs resulted in different decision rates in subsets of the number sort rule, showing how expert performance could be achieved by different expert usage strategies.

**Supplementary Figure 2.**
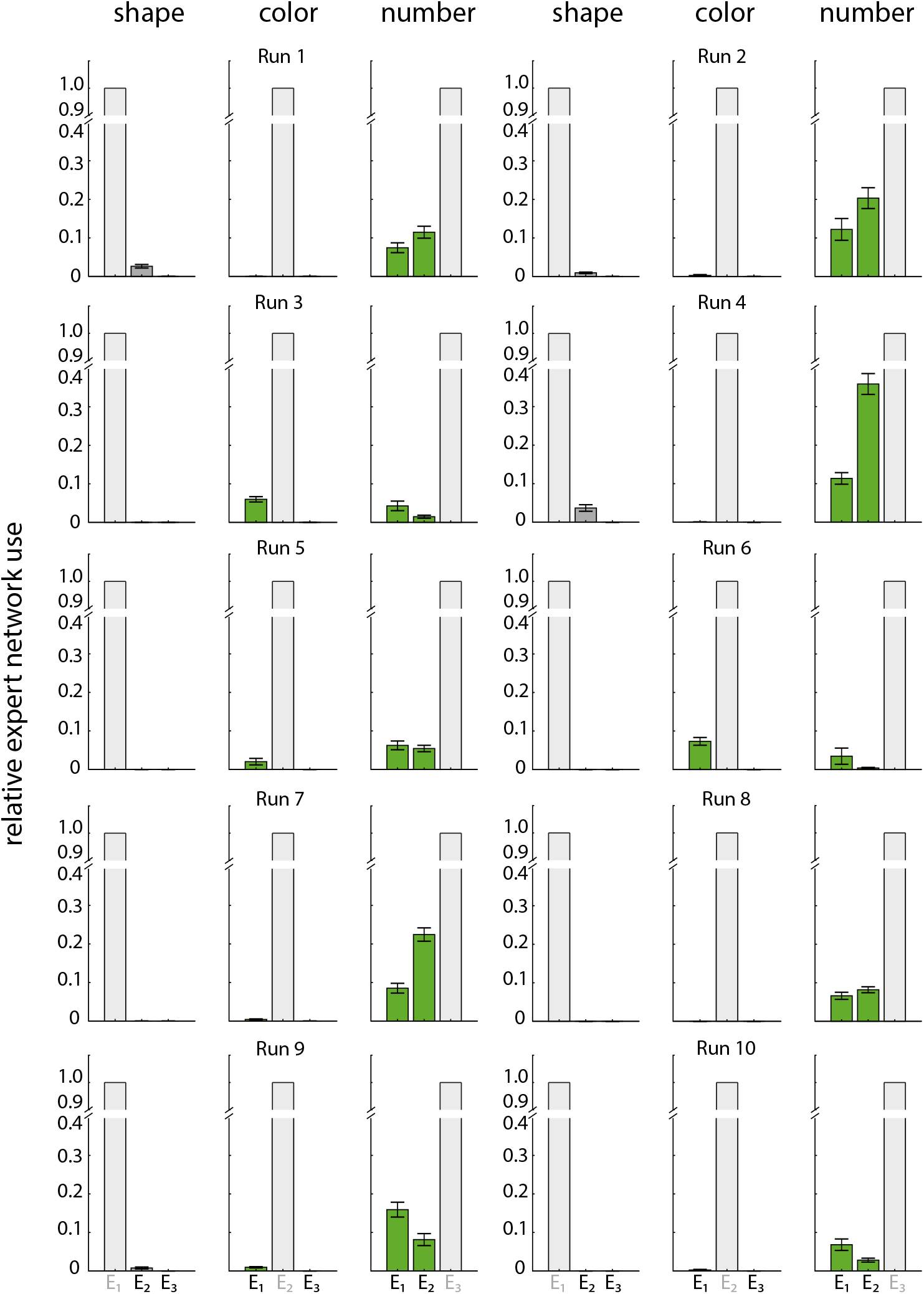
Relative expert network usage in each sort rule for 10 independently trained DynaMoE networks. For each network, after sequential training, the gating network was retuned for each sort rule until expert performance. Weights were then fixed and 1,000 test sorts were done in the same sort rule. Proportion of cards allocated to each expert were normalized to the dominant expert in that sort rule and plotted (see Methods). Color schemes and notations are the same as Fig. 4. Error bars show standard deviation over 1,000 test trials. Independent training runs led to variable levels of expert network usage, while clearly showing a consistent pattern of using experts added before initial training (green bars) more than those added after (grey bars) (Fig. 4).

**Supplementary Figure 4.**
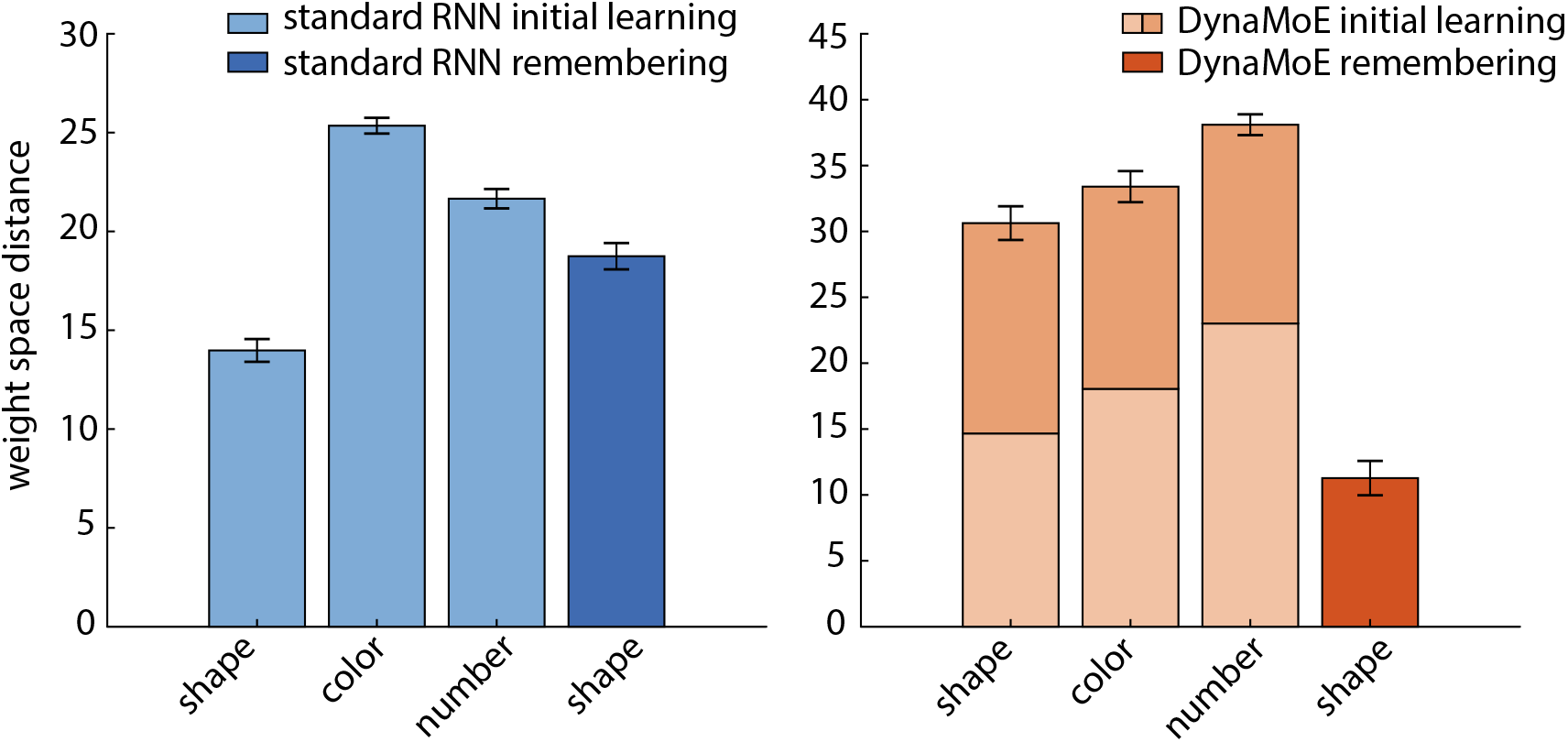
Comparison of networks during initial learning and relearning of a previously learned task in weight space. Bar graphs show movement of networks from Fig. 5 in weight space during sequential training of sort rules in WCST (initial learning of shape rule is leftmost bar of each bar graph). For initial learning of each rule (light blue and light and medium orange bars), networks were trained on 1,250 sorting episodes to allow full convergence to optimal performance (see Fig. 2 and Supplementary Fig. 1). For relearning (dark bars), networks were trained until they reached expert performance criteria (see Methods). Left, Standard RNN. Distances moved in weight space during initial learning of each rule (light blue) and “remembering” of the shape rule (dark blue) are similar. Right, DynaMoE network. In stacked bars, lightest orange indicates stage 1 of training for a task (gating network only), medium orange indicates stage 2 of training (gating and new expert network). Distance moved in weight space during initial learning of each rule is similar. Distance moved during “remembering” shape rule (dark orange) is much less than initial learning of each of the rules (stacked bars). Error bars are SEM for 10 independent training runs of each type of network.

**Supplementary Figure 5.**
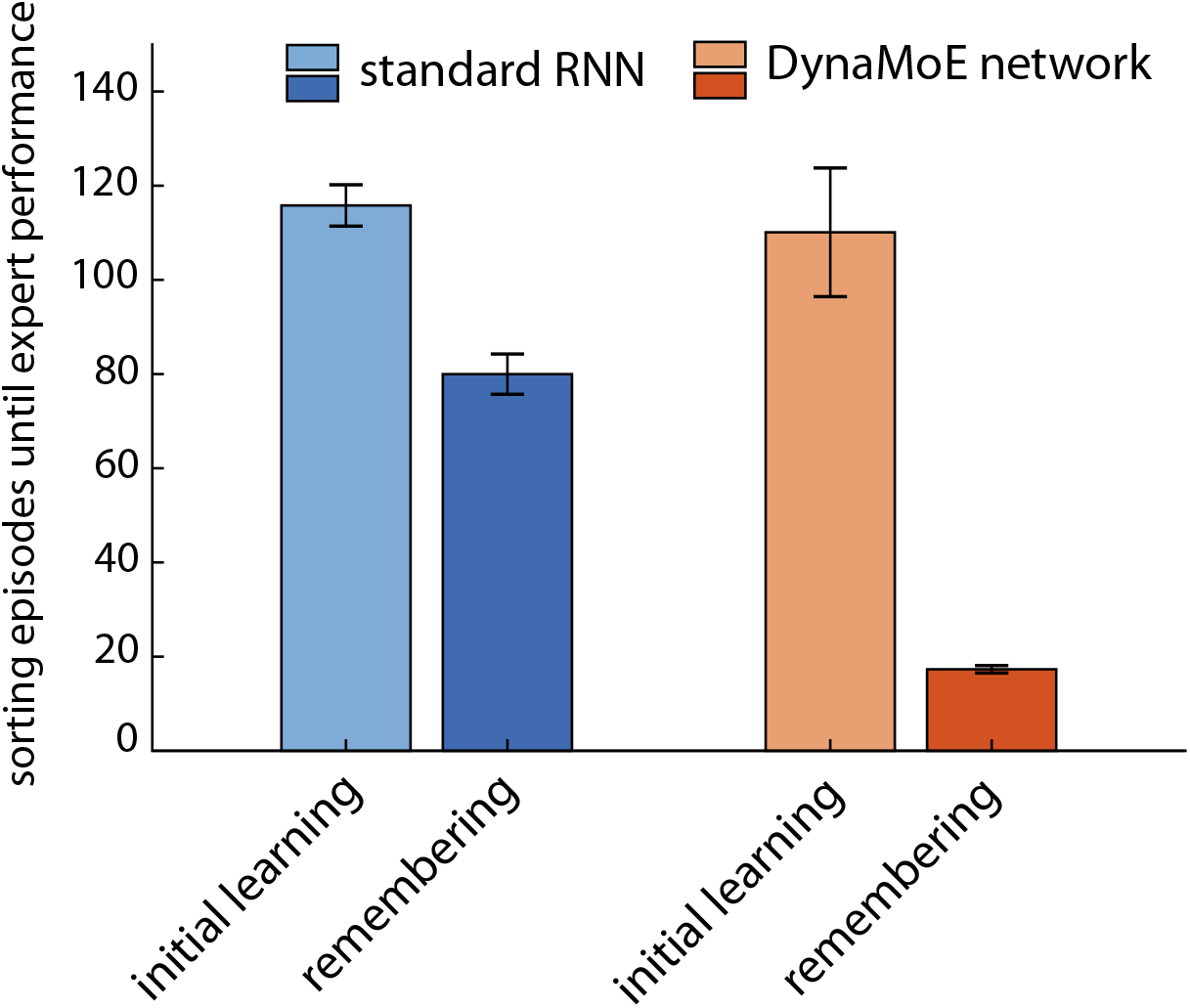
Comparison of number of sorting episodes required to attain expert performance during initial learning compared to during remembering of shape rule after sequential training. For a standard RNN, 31% fewer sorting episodes (*p* = 1.54*e* — 05) were required when remembering compared to initially learning a rule, while for the DynaMoE network 84% fewer sorting episodes (*p* = 2.37*e* — 06) were required for remembering compared to initially learning a rule. Number of sorting episodes required to initially learn the rule were not significantly different between the standard RNN and the DynaMoE network (light colored bars; *p* = 0.70). Error bars are SEM over 10 independent simulations for each condition.

**Supplementary Figure 6.**
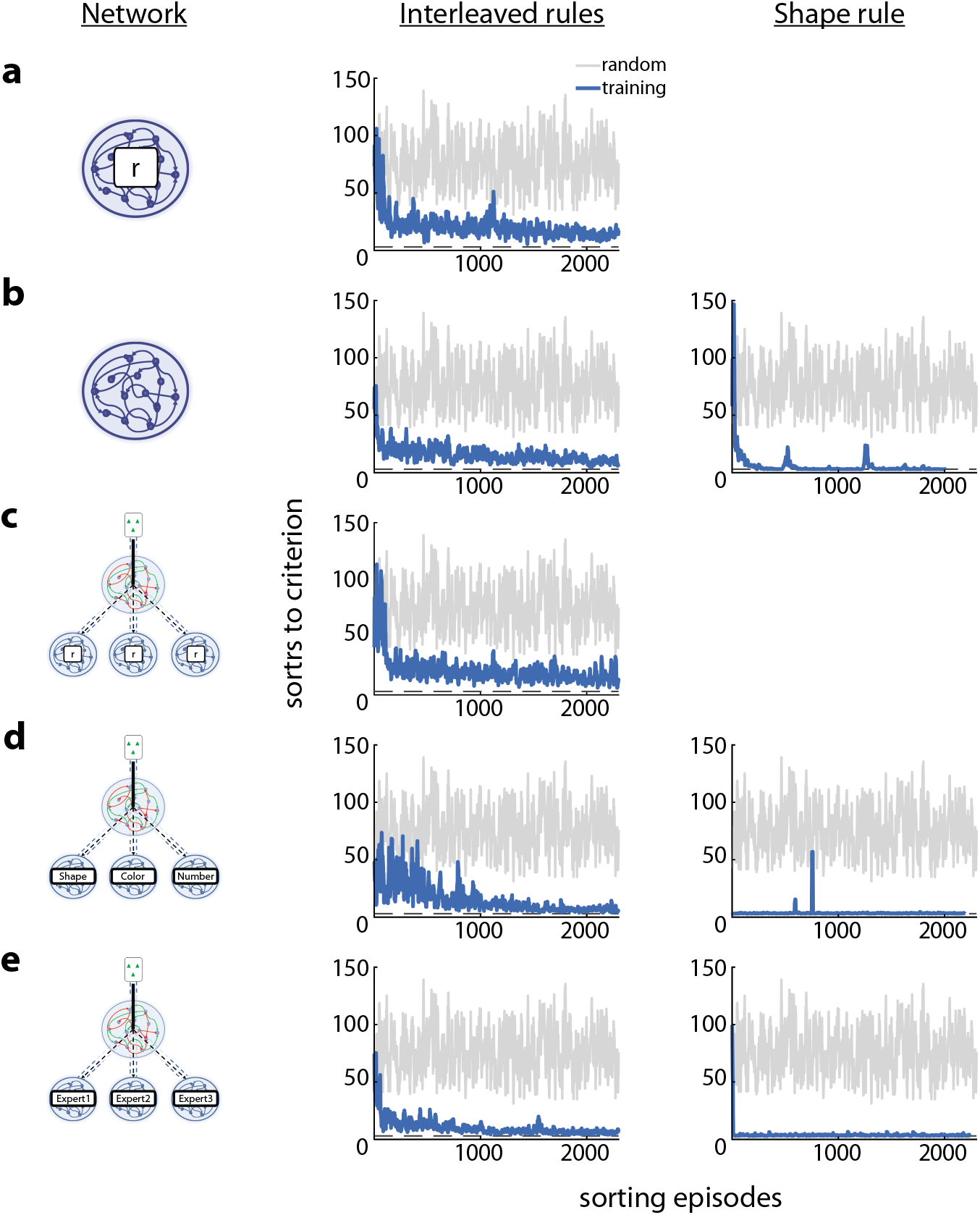
Performance of standard RNN (**a–b**), MoE (**c–d**), and DynaMoE networks (**e**) on the classic interleaved rules WCST and on a single rule after random initialization or pretraining. The first column depicts the network type, the second column shows performance on the classic interleaved rule scenario, and the third column shows performance on the shape rule for networks with pretraining. **a**, A randomly initialized standard RNN rapidly learns the interleaved WCST (mean sorts to criterion (STC) over 10 sorting episodes (SrtEp) <50 by about 50 SrtEp of training). **b**, A standard RNN pretrained sequentially on the shape, then color, then number rules (see Methods) learns the interleaved WCST faster (mean STC <50 by SrtEp 21) and exhibits catastrophic forgetting in the single previously experienced rule. **c–e**, Networks from 1c–d. **c**, MoE network with 3 randomly initialized expert networks (mean STC <50 by SrtEp 109). **d**, MoE network seeded with pretrained rule experts (mean STC <50 from the first SrtEp). Within a single training SrtEp, this network performs near perfect on the single rule scenario. **e**, DynaMoE network trained sequentially in a manner similar to Fig. 2c (see Methods) (mean STC <50 by SrtEp 21) converges to the best performance. It more rapidly achieves near perfect performance on the single rule scenario than the standard RNN, due to its robust memory savings. All simulations were done for 2,500 SrtEp to show convergent learning behavior. In all plots, blue traces are from networks during training and grey traces are random behavior for reference.

**Supplementary Figure 7.**
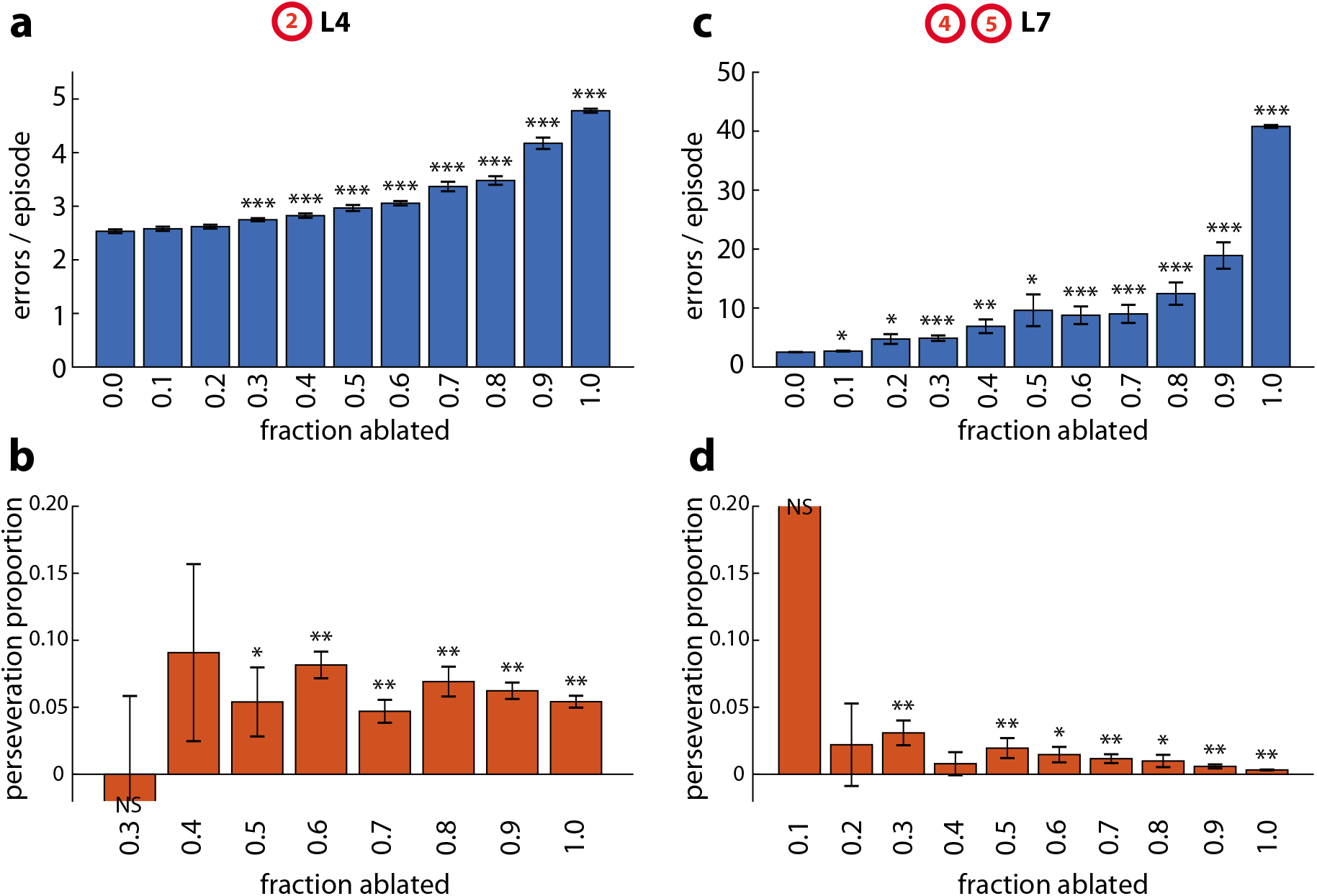
Examples of full spectrum of severity for specific lesions from Fig. 6. **a–b**, L4 - lesions of the internal network dynamics (synaptic connections to the “forget” gate in the LSTM) of DynaMoE’s gating network (region 2 in Fig. 6) **a**, Average total errors per episode after L4 lesions of varying severity. **b**, Average perseveration proportion for lesions that caused significant increase in average total error. **c–d**, L7 - lesions of the synaptic connections to action and value units (region 4 and 5 in Fig. 6). **c**, Average total errors per episode after L7 lesions of vary severity. **d**, Average perseveration proportion for lesions that caused significant increase in average total error. Asterixes indicating significance follow same key as Fig. 6. For all plots, error bars are SEM.

**Supplementary Table 1.** DynaMoE lesion-induced impairments on the WCST. Lesions of the gating network of DynaMoE qualitatively separated into 3 categories based on error rate and perseveration proportion (Fig. 6): (red) lesions that caused significant increase in error rate with small or not significant perseveration proportion; (blue) lesions that caused small but significant increase in error rate with large perseveration proportion; and (grey) lesions that did not significantly change the error rate. The lower end of all ranges is 50% impairment and the high end is 90% impairment for the given lesion type. Lesions labels are same as Fig. 6.

| lesion | 1 |  |  | 2 | 3 | 4 |  | 5 |
| --- | --- | --- | --- | --- | --- | --- | --- | --- |
|  | L1 | L2 | L3 | L4 | L5 | L6 | L7 | L8 |
| error increase (fold) | 7.41 | 1.72 | 10.51 | 1.18 - 1.65 | 2.91 - 7.03 | 2.47 - 6.25 | 3.81 - 7.50 | NS |
| perseveration proportion (%) | 0.29 | NS | NS | 6.67 - 8.22 | 1.11 - 1.51 | 0.81 - 1.49 | 0.59 - 1.95 | NA |

